# Population-Level Activity Dissociates Preparatory Overt from Covert Attention

**DOI:** 10.1101/2025.05.07.652629

**Authors:** Damian Koevoet, Vicky Voet, Henry M. Jones, Edward Awh, Christoph Strauch, Stefan Van der Stigchel

## Abstract

The neural signatures of preparing overt eye movements and directing covert spatial attention overlap as they recruit the same brain areas. Yet, these neural signatures are dissociable at the single-cell level: Specific cells within visuo-oculomotor areas are exclusively involved in motor preparation or covert attention. Nevertheless, it has been proposed that many cells in visuo-oculomotor areas are involved in both motor preparation and covert attention, and consequently their neural signatures should functionally overlap to a large degree. Here, we directly tested this proposal: we combined human (both sexes) EEG with sensitive decoding techniques to investigate whether the neural signatures of preparatory overt and covert attention are dissociable across large-scale neuronal populations. We found that neural decoding reliably discerned whether overt or covert attention was shifted well before saccade initiation. Further, inverted encoding modeling revealed earlier and sharper spatially-tuned activity in preparatory overt than in covert attention. We then asked whether preparatory overt attention achieved sharper spatially-tuned activity by using ‘more-of-the-same’ covert attention, or by recruiting additional spatially selective neural processing. Cross-decoding results demonstrated that preparatory overt attention recruited at least one additional, frontal process. This additional spatially selective process emerged early and likely reflects motor preparation or predictive remapping. To summarize, we found that the neural signatures of overt and covert attention overlap, yet diverge rapidly, in part because overt attention employs an additional spatially selective neural process. Extending beyond a dissociation on the single-cell level, our findings demonstrate that population-level neural activity dissociates preparatory overt from covert attention.

**Significance statement:** The world provides much more visual input than the brain can process simultaneously. Spatial visual attention allows for the selective processing of only the most important parts of this input. Spatial attention shifts either overtly (with an eye movement) or covertly (without an accompanying eye movement). The neural signatures underlying these types of spatial attention have long thought to overlap to a large degree. This strong overlap was thought to only break down at the level of single neurons. In our EEG study, we instead demonstrate that large populations of neurons dissociate overt from covert attention. Our results show that the neural signatures of overt and covert attention differ on a much broader scale than previously thought.

## Introduction

Spatial attention is shifted overtly with, or covertly without an accompanying eye movement (M. Carrasco, 2011; Helmholtz, 1866; Posner, 1980). Preparatory overt^1^ and covert attention are tightly coupled (Rizzolatti et al., 1987, 1994). For example, saccade preparation leads to an attentional shift to the target prior to eye movement initiation (Deubel & Schneider, 1996; Kowler et al., 1995) and saccade trajectories deviate from attended locations (Sheliga et al., 1994). While such behavioral results sparked tremendous interest in the neural signatures of preparatory overt and covert attention, it currently remains unclear to what degree these signatures overlap.

We here define neural overlap as the number of cells coding for multiple functions relative to the number of cells coding exclusively for a single function. To illustrate, consider the following two scenarios wherein a brain contains only ten neurons. In scenario A, 5 neurons exclusively code for motor preparation and 5 neurons would exclusively code for covert attention, there would be no overlap between neural signatures. In scenario B, 1 neuron exclusively codes for motor preparation, 1 other neuron exclusively codes for covert attention while 8 other neurons code for both. The latter scenario implies substantial neural overlap between functions, even despite dissociations at the single-unit level (Awh et al., 2006).

On the one hand, preparatory overt and covert attention seem to neurally overlap to a large degree. For example, they recruit similar (sub)cortical areas including frontal eye fields (FEF) and superior colliculi (Bruce et al., 1985; Corbetta & Shulman, 2002; de Haan et al., 2008; Nobre et al., 2000; Perry & Zeki, 2000). Zooming into the single-cell level, subthreshold microstimulation of FEF cells involved in saccade generation (which would trigger saccades if stimulated above threshold) leads to covert attentional selection (Moore & Armstrong, 2003; Moore & Fallah, 2001). This demonstrates a tight neural link between preparatory overt and covert attention (although neighboring cells were possibly also stimulated, Tehovnik, 1996). On the other hand, it has become increasingly clear that preparatory overt and covert attention are dissociable at the single-cell level: Specific cells code exclusively for motor preparation or covert attention (Juan et al., 2008; Messinger et al., 2021; Schall, 2004). Reconciling these data, Awh et al. (2006) proposed that a large proportion of cells in visuo-oculomotor areas are sensitive to both motor preparation and covert attention, and that therefore preparatory overt and covert attention overlap to a large degree - even despite dissociations at the single-cell level (consistent with scenario B).

If preparatory overt and covert attention indeed neurally overlap to a large degree, dissociations should occur at the single-cell, but not at the population level. By contrast, past work implies that population-level neural activity may dissociate preparatory overt from covert attention. Recently, transcranial magnetic stimulation (TMS) - which affects a large population of neurons - of FEF had differential perceptual effects during preparatory overt and covert attentional selection. Specifically, TMS applied to FEF ameliorated perceptual benefits at validly cued and costs at invalidly cued locations in voluntary covert attention (Fernández et al., 2023). However, TMS at FEF reduced costs at invalidly cued locations but did not mediate benefits at validly cued locations in preparatory overt attention (Hanning et al., 2023). Another example is that pupil size - indicative of activity in attentional networks (Joshi & Gold, 2020; Strauch et al., 2022) - is larger during preparatory overt compared with covert attentional shifts (Koevoet et al., 2023). Despite these tentative findings indicating larger-scale neural dissociations between preparatory overt and covert attention, direct evidence for a neural dissociation at the population level is currently lacking.

We here combined EEG - which reflects synchronous activity of large-scale neuronal populations (Cohen, 2014; Lopes da Silva, 2013; Nunez, 2006) - with sensitive decoding techniques (Brouwer & Heeger, 2009; Grootswagers et al., 2017; Sprague et al., 2018). This allowed us to determine if, when and how preparatory overt and covert attention may dissociate on the population level.

## Materials and Methods

All data and analysis scripts necessary to reproduce the reported results are available via the Open Science Framework: https://osf.io/kd4qp/.

### Participants

We set our target sample to 22 participants based on previous EEG decoding work (e.g. Bae & Luck, 2019; Duncan et al., 2023; Foster, Sutterer, et al., 2017; Foster et al., 2020; Jones et al., 2024). A total of 28 participants took part in the experiment. We excluded three participants due to poor task performance (<50% accuracy in at least one condition) and two other participants because eye-tracking data were unavailable due to technical errors. Our final sample consisted of 23 participants (*M*_*age*_ = 23.22, range: [19-27], 15 female, 8 male) with normal or corrected-to-normal vision and without a diagnosis of, or history with, attention-deficit (hyperactivity-)disorder or autism spectrum disorder. We overshot our target sample size by one participant due to performing artifact rejection and task performance checks in batches. Participants were compensated with course credits or received€8 per hour. The experimental procedure was approved by the Utrecht University ethical review board (approval code: 24-0197).

### Data acquisition and stimuli

Gaze data were recorded from the right eye at 1000 Hz using an EyeLink 1000+ (SR Research, Mississauga, Ontario, Canada). Participants were positioned in a chinrest 67.5 cm from the monitor (ASUS ROG PG278Q; 2560 x 1440; 100 Hz). We calibrated the eye-tracker using a 9-point procedure, and recalibration was performed whenever necessary between blocks. Stimuli were presented using PsychToolbox in MATLAB (Brainard, 1997; Pelli, 1997).

EEG was recorded at 2048 Hz from 30 active electrodes using a BioSemi ActiveTwo system (BioSemi, Amsterdam, The Netherlands). Specifically, we recorded from International 10/20 sites: Fp1, AF3, F7, F3, FC1, FC5, C3, CP1, CP5, P7, P3, Pz, PO3, O1, Oz, O2, PO4, P4, P8, CP6, CP2, C4, FC6, FC2, F4, F8, AF4, Fp2, Fz, and Cz. We additionally placed electrodes ∼1 cm from the outer canthi of the left and right eyes to record horizontal EOG, and placed electrodes above and below the right eye to monitor vertical EOG. We also placed electrodes on the left and right mastoid bones that were used for later offline re-referencing.

Stimuli were displayed on a gray background. The central fixation stimulus was a slightly darker grey dot (0.15°). Cues consisted of blue (17.8 cd/m^2^) and red (15.8 cd/m^2^; counterbalanced) bars (0.12° x 0.25°) placed around the fixation point. Arrays were presented at 4° eccentricity in an invisible circle, and consisted of a single diamond target (1.6° x 1.6°; with a gap on the left or right) and square distractors (1.6° x 1.6°). All stimuli in the array were equidistant (steps of 45°).

### Procedure

Participants performed a cued visual selection task (based on Foster et al., 2020) and identified whether the target diamond stimulus had a missing gap on its left or right side (Figure 1A). Trials started with a brief fixation period (500-600 ms). Then a cue (100% valid) was presented that indicated the upcoming target location. The post-cue delay period was variable (600-900 ms) to discourage participants from making premature saccades toward the target stimulus (following Deubel & Schneider, 1996; Koevoet et al., 2023). A search array was briefly presented (150 ms) and followed by a mask array (400 ms). Participants then indicated whether the missing gap was on the left (‘z’ key) or right (‘/’ key) of the target diamond stimulus. Blank inter-trial intervals ranged between 800 and 1100 ms. Participants were instructed to only blink during the blank inter-trial interval.

**Figure 1:**
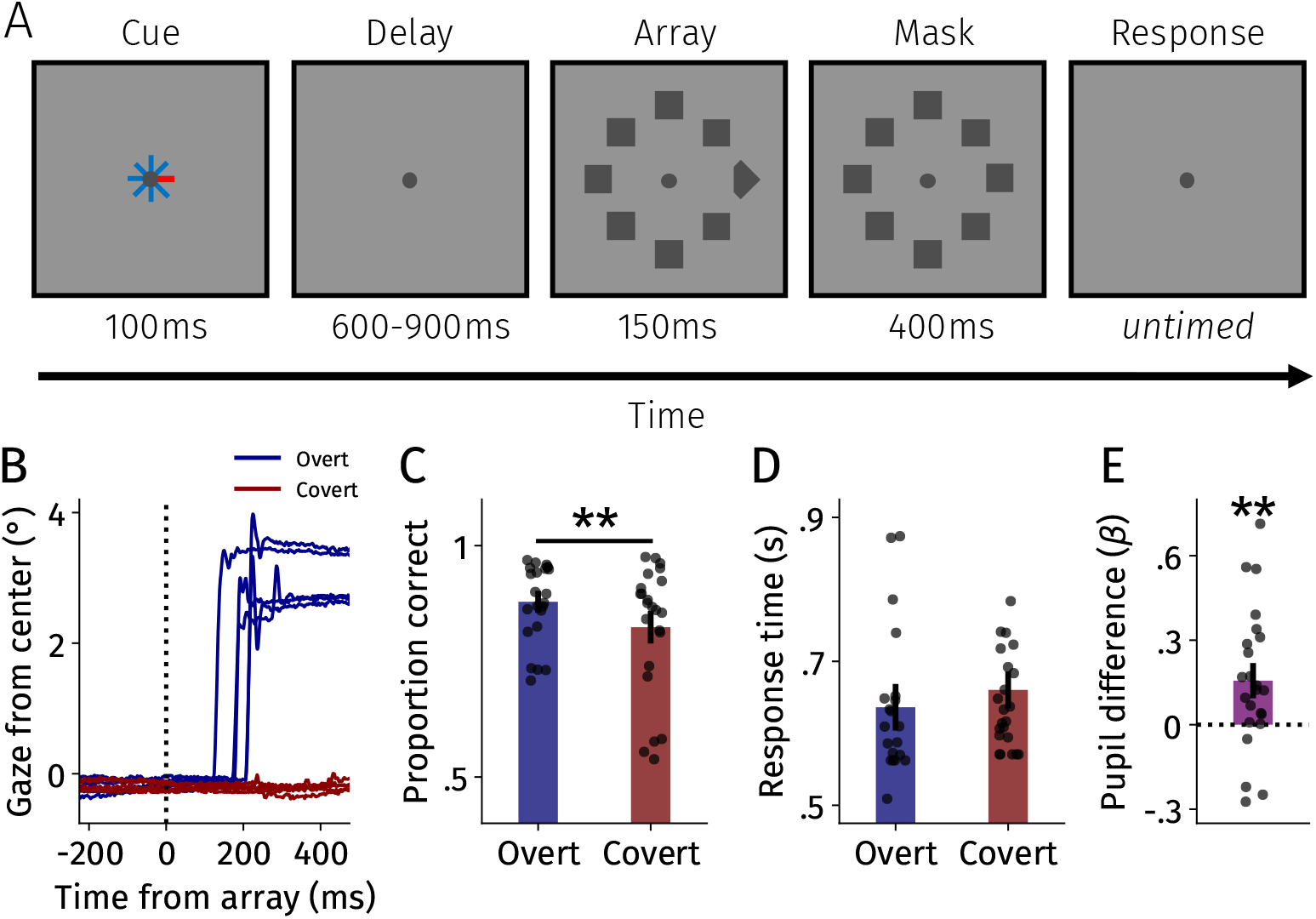
Task and behavioral results. **A** Participants performed a cued visual selection task while shifting overt or covert attention. Participants indicated whether a gap was missing from the left (‘z’) or right (‘/’) part of the target diamond. Cues (100% valid) indicated the target location. **B** Five example gaze traces from a single participant for each condition upon array onset. As is evident from the figure, participants made a saccade toward the target location upon array onset in overt blocks. In covert blocks, participants maintained central fixation throughout the trial. **C** Gap identification accuracy per condition. **D** Response times per condition. **E** Pupil size difference between shift type conditions while controlling for known covariates of pupil size. Higher values indicate a larger pupil size in the overt compared with the covert attention condition. Error bars reflect standard errors of the mean across participants. Black points indicate individual participants. ^**^*p* < .01.

Whereas in the covert condition, participants were instructed to maintain central fixation throughout the trial, in the preparatory overt condition, participants were instructed to make a saccade as fast as possible to the target location upon array onset (see Figure 1B). Participants were given feedback based on their gaze behavior. If gaze deviated more than 2° from central fixation in the covert condition, participants were shown “Keep still!”. In the overt condition, participants received feedback when they made premature (i.e. prior to array onset; “Too early!”) or slow saccades (>400 ms; “Too slow!”).

Each block consisted of 64 trials and blocks of preparatory overt and covert trials alternated. We counterbalanced across participants whether they would start with the overt or covert condition. Participants completed at least 16 blocks but could choose to complete more until a maximum of 20 blocks was reached. Prior to the experimental blocks, participants completed one block of practice trials per condition to get familiar with the experiment. Participants were free to take breaks between blocks. Participants saw their mean accuracy and response time performance in their last completed block during each break.

### Data processing and artifact rejection

Data processing and analyses were performed using custom Python scripts (based on Foster, Sutterer, et al., 2017; Foster et al., 2020; Jones et al., 2024; Thyer et al., 2022). Data were re-referenced to the average of the left and right mastoids, epoched around cue onset (−250 - 700 ms), downsampled to 128 Hz and bandpass filtered (0.5-30 Hz). In the selected epoch, no saccades should be present in the data in both the overt and covert conditions. Therefore, neural activity in this epoch can be compared between conditions without ocular artifacts.

We used an automated artifact detection algorithm based on Foster et al. (2020). Eye-tracking data were used to detect trials with ocular artifacts (e.g. saccades or blinks). Gaze data were drift-corrected by subtracting the median gaze position 100-0 ms prior to cue onset for each trial. We took two steps to detect trials with saccades. First, we discarded trials wherein drift-corrected gaze deviated more than 1° from central fixation. Second, we used a 60 ms sliding window of mean gaze that advanced in steps of 10 ms. If at any point in the trial the difference in gaze between the first and last halve of the sliding window exceeded a threshold of 0.5°, we excluded the trial. All trials wherein the pupil size data were missing at any point were discarded to omit trials with blinks.

Next, the algorithm checked the EEG voltage data itself for potential artifacts. Trials with voltage drifts (i.e. skin potentials) were identified by calculating the absolute voltage difference between the first and last quarter of the trial. We discarded trials if the difference exceeded 40 *µ*V. High-frequency noise (i.e. muscle artifacts) was identified by calculating peak-to-peak amplitudes in ∼15 ms windows (i.e. two samples at 128 Hz) advanced in ∼15 ms steps across the entire trial. Trials were determined to contain high-frequency noise if peak-to-peak amplitudes exceeded 120 *µ*V, and were discarded. We checked for sudden steps in voltage changes by calculating peak-to-peak amplitudes in a window of 250 ms that advanced in steps of ∼21 ms. We examined peak-to-peak amplitudes obtained with the sliding window above, and if the amplitude exceeded 60 *µ*V at any point, the trial was discarded. Lastly, we discarded trials if absolute voltage values exceeded 80 *µ*V at any time during the trial (Thyer et al., 2022). An average of 990.09 trials (*SD* = 127.16) were included per participant in the final analyses.

Data were baseline corrected by subtracting the median voltage within 250 ms prior to cue onset in each trial for each electrode separately. These baseline-corrected data served as the input for further neural analyses.

### Multivariate classification analysis

We conducted a multivariate classification analysis to discern between preparatory overt and covert attention (based on Adam et al., 2020; Thyer et al., 2022). To this end, we trained logistic regression models to classify trials as preparatory overt or covert based on baseline-corrected voltage data (C. D. Carrasco et al., 2024; Grootswagers et al., 2017). Although classification was reliable even at the single trial level, we averaged randomly chosen groups of five trials to increase power (Adam et al., 2020; Grootswagers et al., 2017). Data were split into training (75%) and test sets (25%), while ensuring an equal number of trials for each location and condition (i.e. cross-validation). Following Thyer et al. (2022) and Jones et al. (2024), we standardized the training data (using *sklearn*.*preprocessing*.*StandardScaler*) and standardized the test data using the mean and standard deviation from the training data (Pedregosa et al., 2011). To evaluate model performance, the trained classifier classified trials as preparatory overt or covert in the held-out test set, as well as in a shuffled held-out test set (i.e. shuffled condition labels). This allows to compare actual classifier accuracy with empirical chance performance. Classification was performed at every time point with 1,000 test-training splits of the data.

### Inverted encoding modeling

To track the spatiotemporal dynamics of preparatory overt and covert attention, we employed inverted encoding modeling (IEM, Figure 2; Brouwer & Heeger, 2009; Sprague et al., 2018). Although IEMs in combination with alpha oscillations sensitively track the allocation of spatial attention (Foster, Sutterer, et al., 2017; Foster et al., 2020; Samaha et al., 2016; van Moorselaar et al., 2018), alpha oscillations in isolation may not be able to pick up on all relevant processes involved in saccade preparation. For example, results from Günseli et al. (2024) indicate that the spatial selectivity of saccade preparation may not be limited to alpha-band activity. Therefore, we here instead focused on analyzing voltage patterns across the scalp (C. D. Carrasco et al., 2024; Grootswagers et al., 2017) as voltage measures have popularly been employed in studying presaccadic processes (Gutteling et al., 2010; Huber-Huber et al., 2016; Parks & Corballis, 2008; Talcott & Gaspelin, 2021; Talcott et al., 2025; van der Lubbe et al., 2006). For completeness, we report results obtained with alpha-band (8-12 Hz) power in Text S1 and Figure S1.

**Figure 2:**
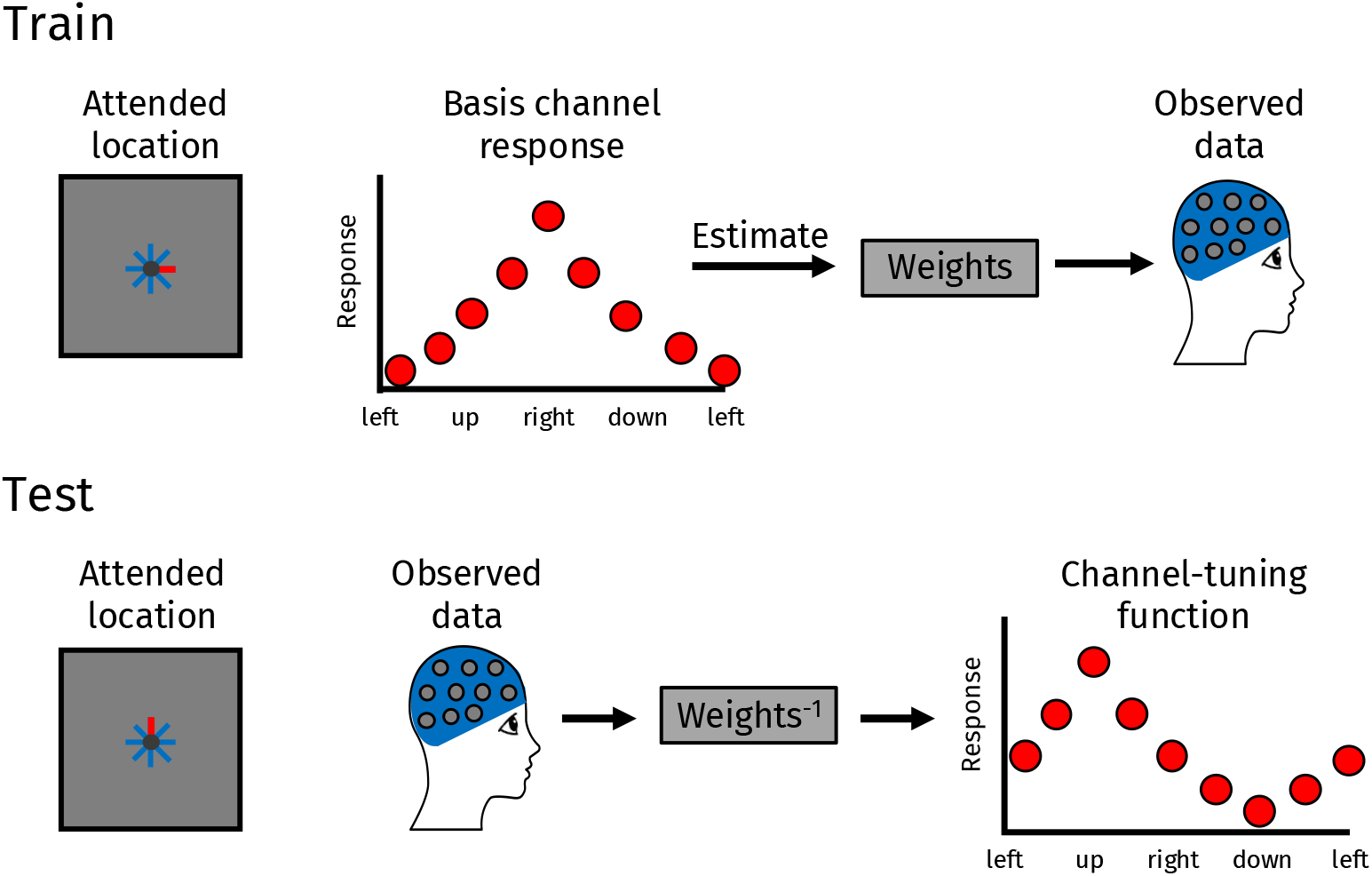
Visualization of the inverted encoding model. Data were first split into train and test sets. Basis channel responses were created for each location separately. Activity was assumed to be highest for the attended location, and followed a half sinusoid (raised to the 7th power) to model the spatial selectivity. To obtain weights, we used these basis channel responses for each location and fit them to the training data. Inverting the obtained weights matrix allows for the reconstruction of channel-tuning functions. Specifically, test data were fit to the inverted weights matrix to reconstruct channel-tuning functions for each location separately.

We reconstructed spatially sensitive channel tuning functions (CTFs) from the voltage topography across time. We assumed that voltage patterns at each electrode reflected the weighted average of eight spatial channels, which were each selective to a specific location. We modeled these basis channel tuning functions using a half sinusoid raised to the 7th power that were centered on all possible target locations (as in Foster, Bsales, et al., 2017; Foster, Sutterer, et al., 2017; van Moorselaar et al., 2018).

For each iteration, data were split into training (2 blocks) and test (1 block) data while an equal number of trials at each cued location as well as for each shift type was ensured per block. Note that for all IEM analyses, unless explicitly mentioned, we trained and tested on data from both shift type conditions. We then fit the basis channel tuning functions to the training data using a general linear model (using *numpy*.*linalg*.*lstsq*), and we obtained weights at each electrode. We then inverted the weights matrix to reconstruct putative spatially sensitive CTFs separately per condition in the test data. The procedure was conducted at each time point (every ∼7.81 ms), and to consider all trials in the analyses we ran 100 iterations per participant over time. We recentered CTFs (i.e. the center corresponds to the cued location) and averaged the reconstructed CTFs across the eight possible locations. This allowed us to track spatial selectivity of neural activity with a high temporal precision.

In keeping with previous work (Foster, Sutterer, et al., 2017; Foster et al., 2020; Jones et al., 2024; van Moorselaar et al., 2018), we quantified spatial selectivity of the reconstructed CTF by calculating its slope. More specifically, we first averaged channel responses equidistant to the target location, and then estimated the slope of the reconstructed CTF using linear regression. A higher CTF slope indicates greater spatial selectivity.

To visualize CTF slope topographies, we used a searchlight approach (Kriegeskorte et al., 2006). We iteratively selected groups of 5 neighbouring electrodes around the scalp and conducted the IEM procedure and calculated the average CTF slope for the time window of 100-700 ms after cue onset (using 25 iterations per electrode group) (similar to Hajonides et al., 2021). We then averaged CTF slopes across all channel-groups that a specific electrode was included in. This allowed us to obtain specific CTF slope contributions for each electrode separately per condition but collapsed across positions.

### Pupillometry

For the pupillometry analysis, we first ran online saccade detection around array onset. We applied a Savitzky-Golay filter (Savitzky & Golay, 1964) and detected saccade onsets using a velocity threshold of 20°/s (Koevoet et al., 2023; Van der Stigchel & de Vries, 2015). For this analysis, we discarded trials with premature, very fast or very slow saccade onsets (<100 ms and >600 ms) in the overt condition, and trials with any detected saccade in the covert condition. (Note that using this subset of trials for the neural decoding analyses yields identical results.) We reconstructed blinks (Mathôt & Vilotijević, 2022), downsampled the data to 100 Hz, linearly interpolated remaining missing data and baseline corrected using the median pupil size 250-0 ms prior to cue onset (Koevoet, Naber, et al., 2024). Following Koevoet, Van Zantwijk, et al. (2025), we then averaged pupil size prior to saccade initiation (−200 - 100 ms around array onset). We z-scored pupil size and conducted multiple linear regression for each participant separately (Wilkinson notation: z-scored pupil size ∼ shift type ^*^ y target position + x target position + oblique + correct). This allowed us to control for individual differences in pupil size as well as to control for effects of known covariates of pupil size, such as task difficulty and shift direction (Beatty, 1982; Kahneman, 1973; Koevoet, Strauch, et al., 2024; Koevoet, Van Zantwijk, et al., 2025; Koevoet et al., 2023). We used a two-tailed one-sample *t*-test to test whether the shift type betas reliably differed from 0, effectively testing the difference in pupil size between preparatory overt and covert attention (Figure 1E).

### Statistical analysis

To analyze behavioral data, we calculated average accuracies and median response times per condition (in artifact-free trials). We compared conditions using two-tailed paired-samples *t*-tests and report Cohen’s *d*_z_ as a measure of effect size (Lakens, 2013).

For the classification analysis, we tested whether classification accuracy was above empirical chance. To this end, we first subtracted decoding accuracies in the shuffled from the non-shuffled data. We then tested whether the difference between these accuracies was reliably above 0. To circumvent the issue of multiple comparisons, we used one-tailed one-sample permutation tests (using *mne*.*stats*.*permutation_cluster_1samp_test*) to identify putative clusters and test their reliability (Maris & Oostenveld, 2007).

As for spatial selectivity, we tested whether CTF slopes were reliably above 0 (i.e. indicative of spatial selectivity) separately for the preparatory overt and covert conditions using one-tailed one-sample permutation tests. To compare the spatial selectivity between preparatory overt and covert attention, we subtracted CTF slopes of the covert from the preparatory overt condition and used a two-tailed one-sample test to assess significance. All cluster-based tests were conducted with *α* = .05 using 10,000 iterations.

We compared CTF slopes between conditions in the cross-decoding IEM searchlight topographies with permutation tests separately for each electrode (5,000 iterations).

## Results

### Behavioral and Pupillometry Results

Twenty-three participants performed a cued visual selection task (Figure 1A; Foster et al., 2020) wherein they shifted overt or covert attention while scalp EEG was recorded. In each trial, a cue (100% valid) indicated where the target diamond would later appear. Participants reported whether the gap was on the left or right side of the target diamond.

The task required participants to attend overtly or covertly in alternating blocks. In the preparatory overt condition, participants were instructed to saccade toward the cued location upon array onset. In the covert condition, participants were asked to maintain central fixation throughout the trial. In Figure 1B, we present five example gaze traces per condition from a single participant for illustration.

We analyzed accuracy and response time as behavioral outcomes. Accuracy was generally high (Figure 1C). Participants were more accurate in the preparatory overt compared with the covert condition (*t*(22) = 3.11, *p* = .005, *d* = 0.65). Response times did not differ significantly between conditions (Figure 1D, *t*(22) = 1.48, *p* = .153, *d* = 0.31). These data indicate that participants were well able to perform the task, and that participants were more accurate in overt compared with covert trials.

We recently demonstrated that preparatory overt attention causes more pupil dilation than shifting covert attention (Koevoet et al., 2023). We here compared pupil size prior to saccade onset (−200 - 100 ms around array onset) (Koevoet, Van Zantwijk, et al., 2025; Koevoet et al., 2023) between shift types while controlling for known covariates of pupil size (e.g., task difficulty (Beatty, 1982; Kahneman, 1973; Koevoet, Strauch, et al., 2024; Mathôt, 2018; Sirois & Brisson, 2014; Strauch et al., 2022; van der Wel & van Steenbergen, 2018) and shift direction (Koevoet, Van Zantwijk, et al., 2025; Koevoet et al., 2023); see Methods). In line with our previous work, we observed that preparatory overt attention caused a larger pupil size prior to saccade initiation compared with covert attention (Figure 1E; one-sample *t*-test: *t*(22) = 2.98, *p* = .007, *d* = .62).^2^ This is a first sign that population-level activity (Joshi & Gold, 2020; Strauch et al., 2022) may differ between shift types.

### Preparatory Overt and Covert Attention Diverge Rapidly

Our main goal was to investigate whether activity in large-scale neuronal populations (as reflected in EEG (Lopes da Silva, 2013; Nunez, 2006)) discerned between preparatory overt and covert attention. Our neural analyses focused on the period preceding array onset because participants did not (yet) initiate a saccade toward the target location in either condition (as in the pupillometry analysis). This allowed for a clean comparison of neural activity between shift types without ocular artifacts. In our first neural analysis, we used time-resolved multivariate classification to decode whether preparatory overt or covert spatial attention was shifted.

We could decode whether participants shifted preparatory overt or covert attention early in the trial. Specifically, we observed significant decoding already from 70 ms after cue onset, which remained reliable throughout the delay period (70-700 ms; *p* < .0001; Figure 3A). This dissociation preceded saccade initiation by at least 630 ms. Therefore, neural processes unfolding well before saccade initiation readily dissociated between preparatory overt and covert attention. Decoding was highly robust: We averaged classification accuracy across the delay period (100-700 ms after cue onset; Figure 3B), and observed above empirical chance classification for each participant (one-sample *t*-test: *t*(22) = 10.06, *p* < .0001, *d* = 2.10). Control analyses demonstrated that none of the decoding results throughout the paper can be ascribed to (premature or miniature) eye movements (Text S2 and Figures S2-3).

**Figure 3:**
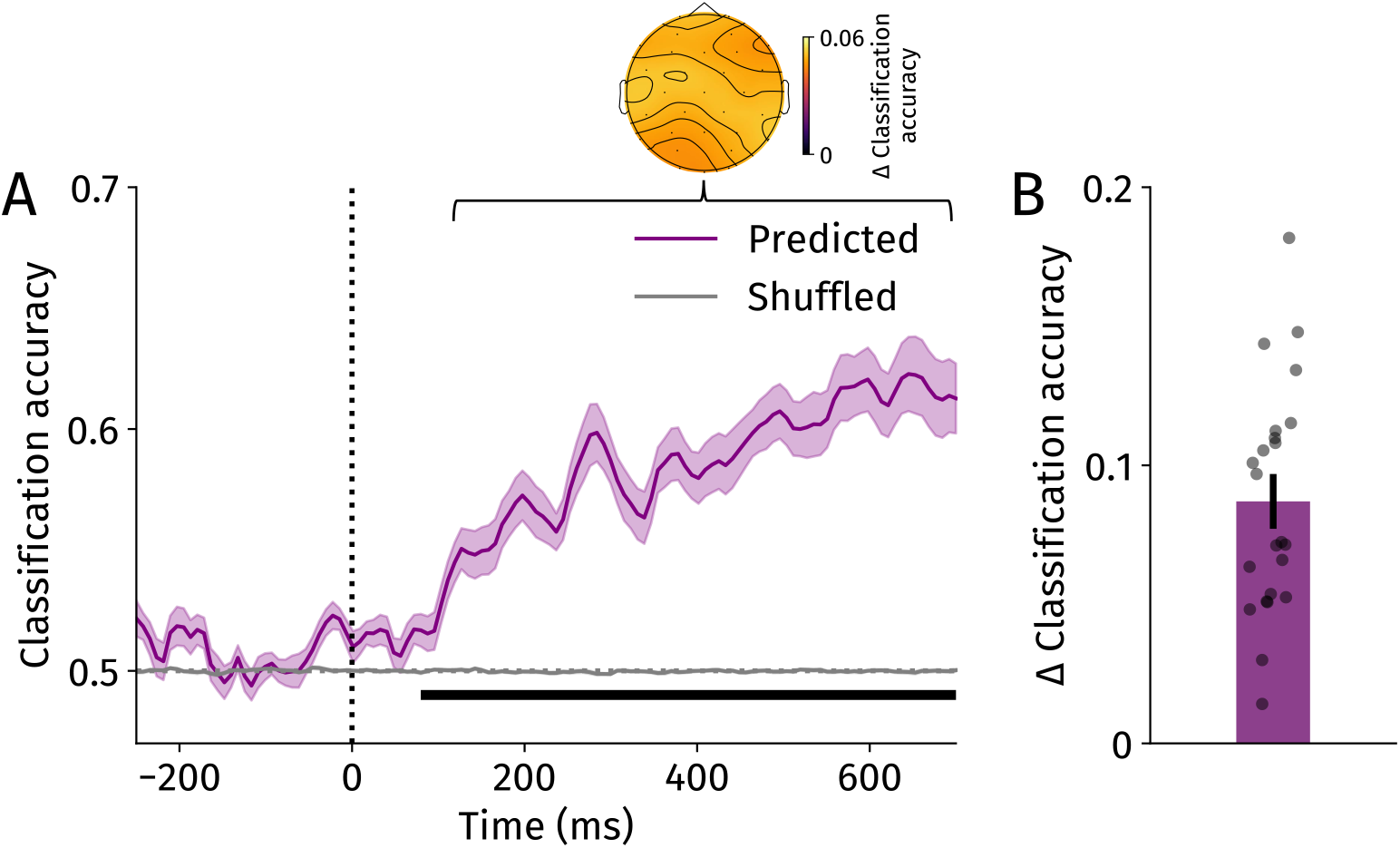
Multivariate classification results. **A** Time-resolved classifiers were trained to identify trials as overt or covert. The purple line represents classification accuracy, while the gray line depicts accuracy on shuffled trial labels (i.e. empirical chance). Shaded error bands reflect standard errors of the mean across participants. Horizontal black line indicates a significant cluster where classifier accuracy was higher than empirical chance (*p* < .0001). Topography reflects the outcome of a searchlight analysis averaged across 100-700 ms after cue onset. **B** The averaged classification accuracy (100-700 ms after cue onset) difference between the actual and shuffled data. Positive values indicate better classification performance in the non-shuffled compared with the shuffled data. Black points represent individual participants. Error bar holds standard error of the mean.

We visualized topographies of classification accuracy using a searchlight analysis (Kriegeskorte et al., 2006). To this end, we iteratively selected groups of 5 neighboring electrodes and repeated the classification analysis in each group, and averaged accuracies (actual - empirical chance) across the delay period (100-700 ms after cue onset). This analysis revealed that neural activity across the scalp discerned whether preparatory overt or covert attention was shifted. Together, the classification results demonstrate that activity in large-scale neuronal populations discerned between preparatory overt and covert attention rapidly after cue onset, and well before saccade initiation.

### Sharper Spatial Tuning in Preparatory Overt than in Covert Attention

The above analysis established that preparatory overt and covert attention diverge rapidly after cue onset. But *how* do these shift types differ? Likely candidates that contribute to differences between preparatory overt and covert attention include motor preparation and predictive remapping (Casteau & Smith, 2019; Koevoet et al., 2023; Li, Hanning, & Carrasco, 2021; Smith & Schenk, 2012), which are inherently spatial processes. We therefore focused on how preparatory overt and covert attention may differ in terms of spatially selective neural activity. To this end, we employed inverted encoding modeling (IEM; Brouwer & Heeger, 2009; Sprague et al., 2018), which allows for the spatiotemporal tracking of attention (Foster, Sutterer, et al., 2017; Samaha et al., 2016). Specifically, we employed IEMs to track whether spatial selectivity of large-scale neuronal populations differed between preparatory overt and covert attention (Figure 4).

**Figure 4:**
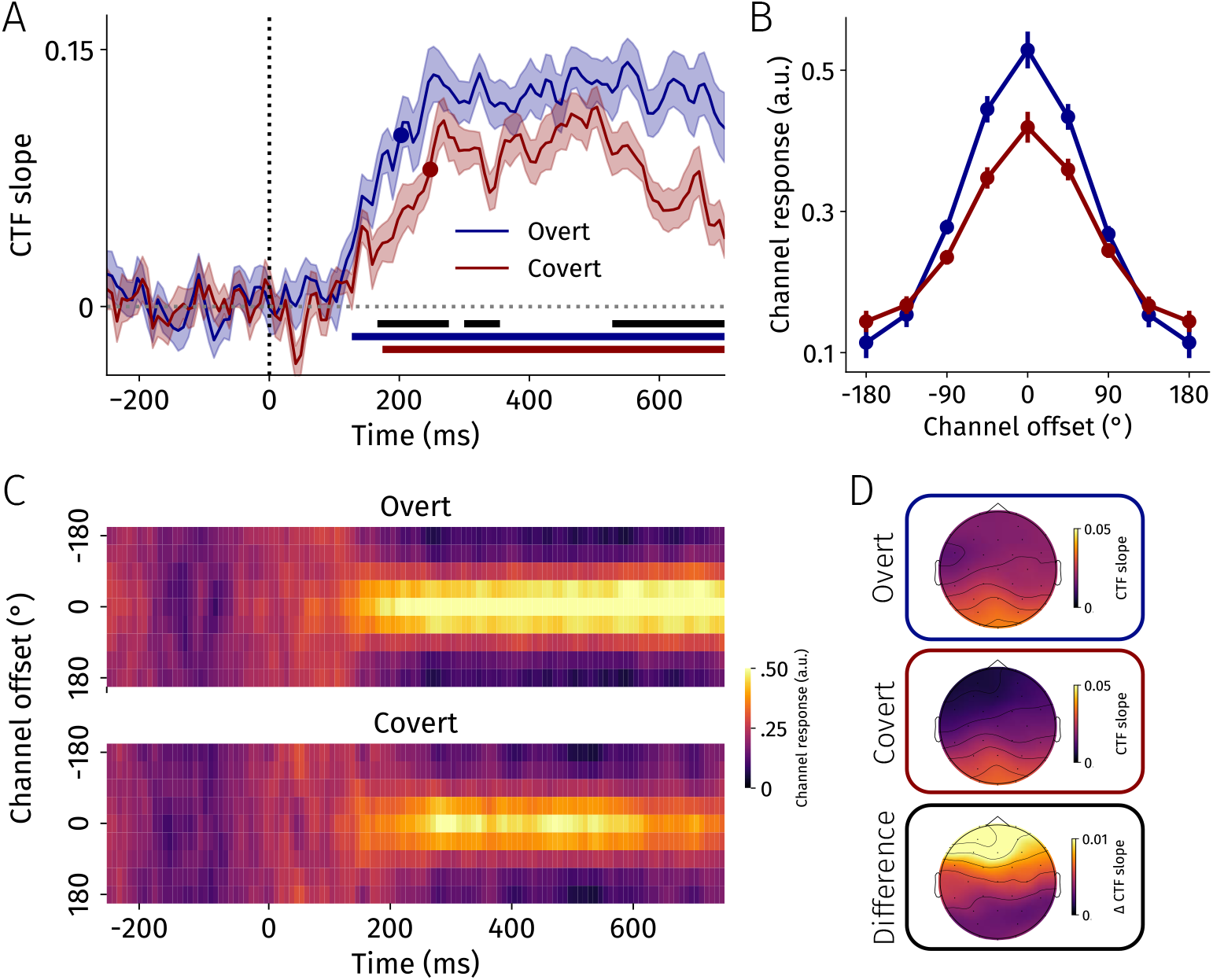
Inverted encoding modeling results. **A** CTF slopes across time for the overt (blue) and covert (red) conditions. The reconstructed CTF’s slope is a measure of spatial selectivity, where higher values represent stronger spatial selectivity. Blue and red circle markers indicate 75%-peak latencies for preparatory overt and covert CTF slopes, respectively. Shaded error bands reflect standard errors of the mean across participants. Blue and red horizontal lines indicate significant CTF slope clusters for preparatory overt and covert data, respectively (*p* < .0001). Horizontal black lines indicate significant clusters between conditions (*p* < .05). **B** Averaged (100-700 ms after cue onset) reconstructed CTFs for the overt (blue) and covert (red) conditions. For both tuning curves, responses are strongest for the cued location (i.e. channel offset of 0) and drop off at further distances. Error bars reflect standard errors of the mean across participants. **C** Heat maps of channel responses across time per shift type. **D** Searchlight-based visualizations of channel contributions to CTF slopes across the scalp. Difference topography was calculated by subtracting the covert from the overt topography; positive values thus correspond to higher slopes in the overt condition (and vice versa).

In line with previous work (e.g. Foster, Sutterer, et al., 2017; Samaha et al., 2016), our results revealed reliable spatial selectivity of neural activity when shifting *covert* attention. Spatial selectivity emerged 164 ms after cue onset and remained throughout the delay (*p* < .0001; Figure 4A,B,C). We here extend this by demonstrating that preparatory *overt* attention also caused spatially-tuned neural activity (117-700 ms, *p* < .0001; Figure 4A,B,C) (also see Günseli et al., 2024). Thus, IEMs precisely tracked spatial attention across time in both conditions.

We then tested whether the spatial selectivity of neural activity differed between shift types. We observed sharper spatially-tuned neural activity in overt compared with covert trials (Figure 4A,B,C; see Figure S4 for individual data). Specifically, we found two early (cluster 1: 144-273 ms, *p* = .002, cluster 2: 289-323 ms, *p* = .015) and a later significant cluster in the delay period (cluster 3: 516-700 ms, *p* < .001). The preparation of an imminent saccade caused sharper spatially selective neural activity compared with directing covert attention already early after cue onset, and thus well before saccade initiation (>550 ms). Additionally, we compared the onset latencies of spatially selective neural activity between conditions. To this end, we used a jackknife resampling procedure (Miller et al., 1998). We estimated the 75%-peak latency across conditions (circle markers in Figure 4A). We observed that spatially selective neural activity emerged significantly earlier in preparatory overt than in covert attention (*t*(22) = 4.04, *p* < .001, *d* = 0.84). These results indicate that preparatory overt and covert attentional shifts diverge early, in part due to earlier and sharper spatial selectivity of large-scale neural activity to a cued location.

We created topographies of spatial selectivity in the preparatory overt and covert conditions using a searchlight approach (Figure 4D; Kriegeskorte et al., 2006). To this end, we iteratively selected groups of 5 neighboring electrodes and repeated the IEM procedure on each group, and averaged CTF slopes across the delay period (100-700 ms after cue onset). This allowed us to obtain the contribution of each electrode to the observed spatial selectivity (i.e. CTF slope) per condition but collapsed across cued locations. As expected, we observed considerable contributions to spatial selectivity in posterior (occipital and parietal) sites for both shift types (van Moorselaar et al., 2018). In addition, frontal channels contributed more to spatial selectivity during preparatory overt compared with covert attentional shifts. Together, the topographies reveal overlapping spatial selectivity in posterior areas across shift types, and a possible spatial selectivity in frontal areas that may only be present during preparatory overt but not covert attention.

We also examined whether neural spatial selectivity predicted behavioral outcomes. We first correlated the difference in CTF slopes between shift types (i.e. overt-covert) with the accuracy difference between shift types, and observed no significant relationship (*r*(21) = .017, *p* = .94). To examine relationships with behavior with increased statistical power, we then turned to within-subjects comparisons. As there were limited incorrect trials, we examined response times. In brief, we conducted a median split on response times, and found that CTF slopes only weakly modulated response times (for details, see Text S3 and Figure S5).

### Preparatory Overt Attention Recruits an Additional Spatially Selective Neural Process

Preparatory overt attention was accompanied by sharper spatially-tuned neural activity tracking the cued location compared with covert attention. Pushing this further, we asked whether this difference was due to overt attention recruiting ‘more-of-the-same’ covert attention, or because overt attention recruits (at least) one additional spatially selective neural process (Figure 5). We previously hypothesized that the larger pupil size in preparatory overt attention (Figure 1E) may be attributed to the recruitment of additional necessary neural processes such as motor preparation and remapping (Koevoet et al., 2023).

**Figure 5:**
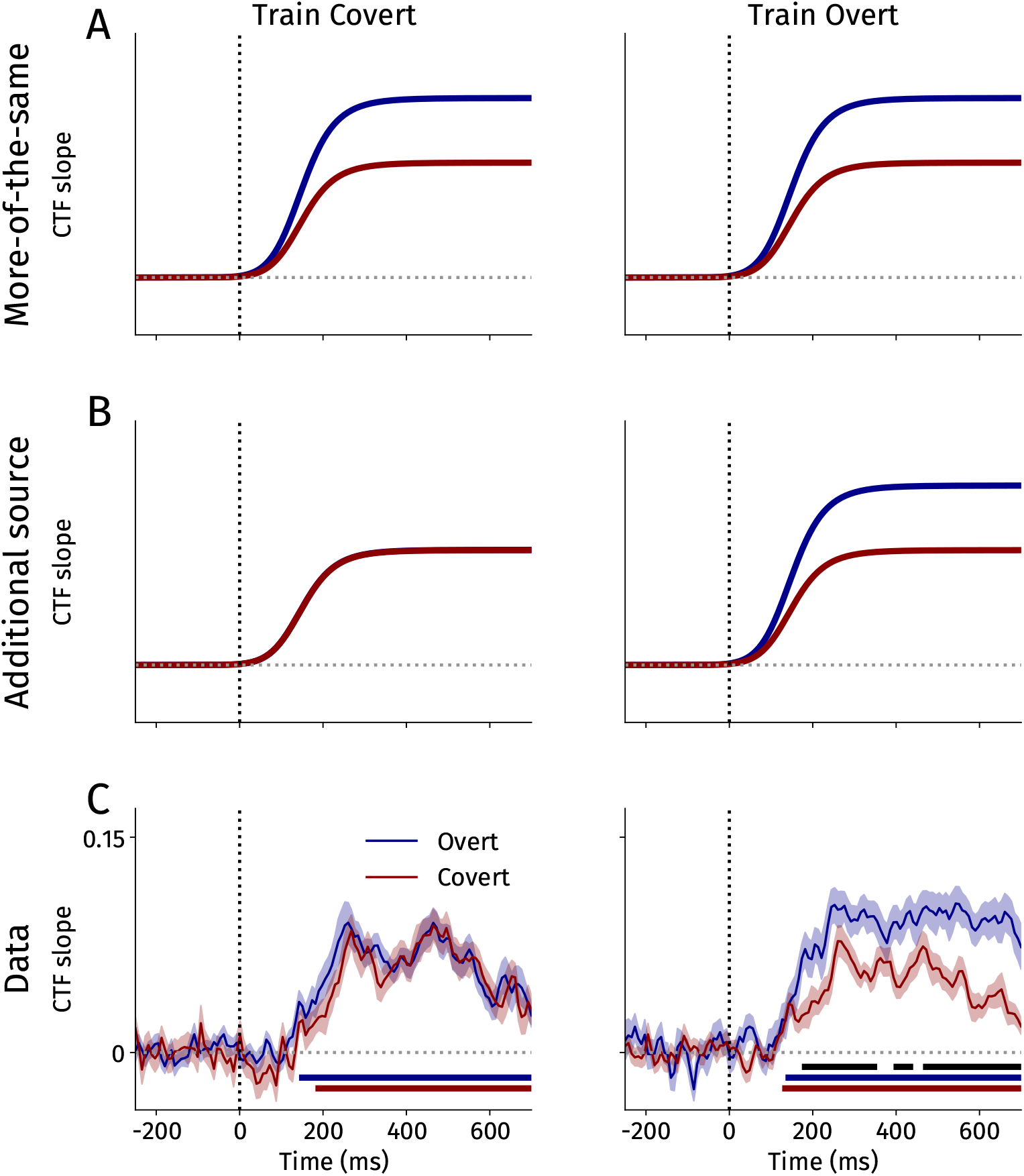
Hypothetical and actual inverted encoding modeling (IEM) results when training on only covert or overt trials. For each graph, we visualized CTF slopes across time for the overt (blue) and covert (red) conditions when training only on covert (left) and overt (right) data. **A** Hypothetical IEM results from a ‘more-of-the-same’ account. **B** Hypothetical IEM results from an ‘additional source’ account. **C** Actual IEM results when training on only covert or overt trials. Shaded error bands reflect standard errors of the mean across participants. Blue and red horizontal lines indicate significant CTF slope clusters for overt and covert data, respectively (*p* < .0001). Horizontal black lines indicate significant clusters between conditions (*p* < .05).

We reasoned that a ‘more-of-the-same’ account would predict that preparatory overt attention makes stronger use of the neural signals present when shifting covert attention (Figure 5A). In contrast, if overt attention employs an additional spatially selective neural process, this process should be absent when shifting covert attention (Figure 5B). To arbitrate between these accounts, we now trained IEMs on only one shift type but tested them on both conditions (i.e. cross-decoding).

When exclusively training on covert data, a ‘more-of-the-same’ account would predict a difference between shift types in terms of neural spatial selectivity (Figure 5A). In contrast, when training exclusively on covert data, the difference in neural spatial selectivity between overt and covert conditions disappeared (all *p*s > .16; Figure 5C). We did observe reliable neural spatial selectivity for the cued location in both conditions, demonstrating overlap in spatially-tuned neural activity between conditions (overt: 133-700 ms, *p* < .0001; covert: 172-700 ms, *p* < .0001). This argues against a more-of-the-same account and instead favors the interpretation that overt attention employs an additional spatially selective neural process.

If so, when training exclusively on overt data - where the IEM would once again have access to the putative additional spatially selective process - the difference in neural spatial selectivity between conditions should reemerge (Figure 5C). We indeed observed sharper spatial tuning in preparatory overt compared with covert attention when training on overt data exclusively (cluster 1: 164-352 ms, *p* = .001; cluster 2: 383-438 ms, *p* = .018; cluster 3: 453-700 ms, *p* < .001). Again, we also found spatially selective neural activity in both conditions separately, indicating overlap in the employed spatially selective neural processes (overt: 125-700 ms, *p* < .0001; covert: 117-700 ms, *p* < .0001).

To formally test whether the higher spatial selectivity in overt compared with covert attention depended on the training data, we calculated the difference in averaged CTF slopes (100-700 ms after cue onset) between shift types when training only on overt or covert data. Comparisons confirmed that the difference between spatial selectivity in overt compared with covert trials is higher when training only on overt compared with training only on covert data (*t*(22) = 4.11, *p* < .001, *d* = 0.86).

Using the same searchlight approach as before (Figure 4D), we visualized topographies of the cross-decoding results (Kriegeskorte et al., 2006). Across all topographies (averaged across 100-700 ms post-cue), posterior electrodes showed the most prominent spatial selectivity to cued target locations–which is in line with past work (e.g. van Moorselaar et al., 2018). Examining the difference topographies, we observed that when exclusively training on preparatory overt trials, frontal electrodes exhibited enhanced spatial selectivity when tested in preparatory overt compared with covert trials (Figure 6; top right). In contrast, when exclusively training on covert trials, we found no differences in spatial selectivity at any electrode between testing conditions (Figure 6; bottom right). This demonstrates that an additional spatially selective process is recruited in frontal cortex during preparatory overt attention that is not present when shifting covert attention.

**Figure 6:**
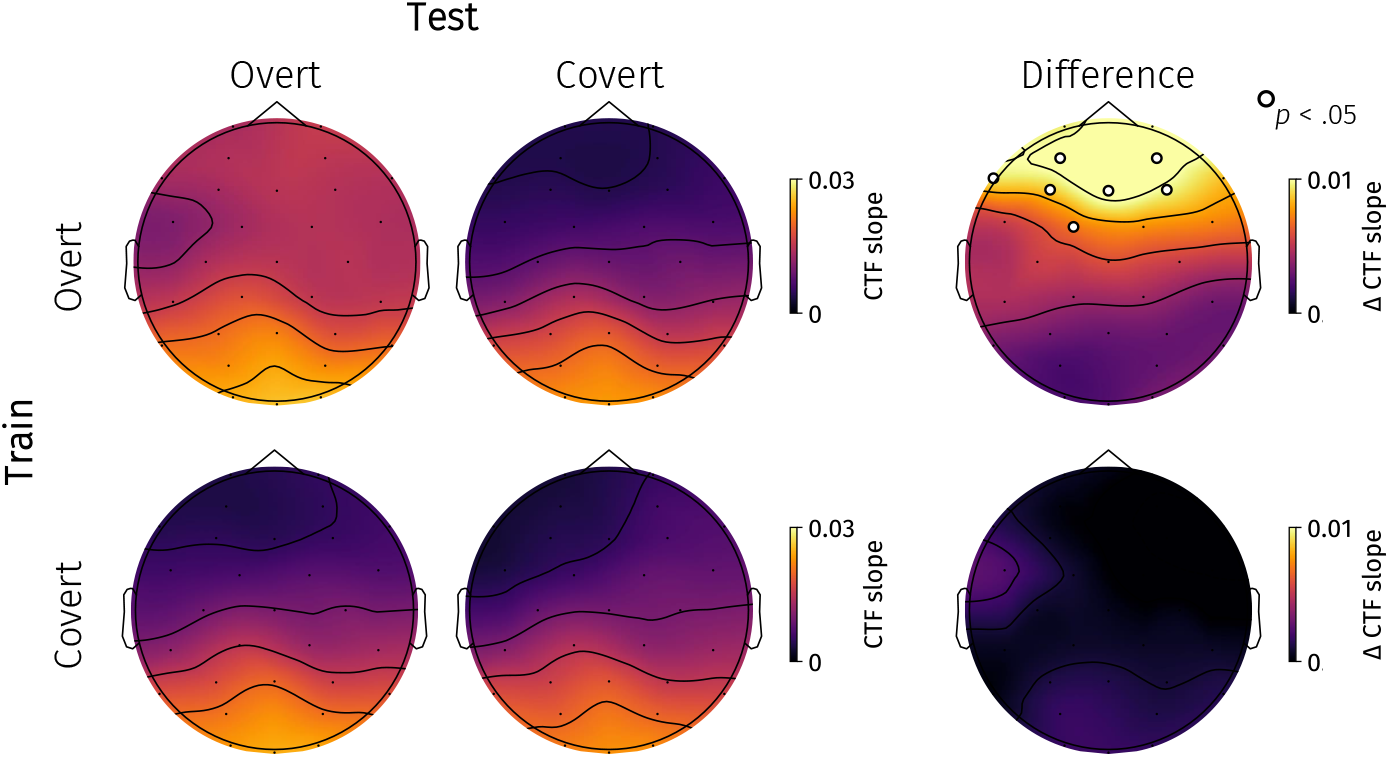
Searchlight results from the cross-decoding IEM analysis. CTF slopes were averaged 100-700 ms after cue onset. The top and bottom rows represent the CTF slopes when exclusively training on preparatory overt and covert trials, respectively. The first and second columns indicate whether the IEMs were tested on overt or covert trials, respectively. The third column represents differences (overt - covert) in spatial selectivity between conditions estimated per electrode. White-black circles indicate electrodes where CTF slopes were significantly steeper when tested on preparatory overt compared with covert trials (*p* < .05).

Past work has suggested that, in some cases, cross-decoding asymmetries may arise from differences in signal-to-noise ratio instead of actual neurocognitive effects (van den Hurk & Op de Beeck, 2019). We examined whether differences in signal-to-noise ratio were likely to explain our cross-decoding results (for details, see Text S4 and Figures S6). To this end, we simulated EEG data based on properties from our observed EEG data at differing signal-to-noise ratios. We found a different pattern of cross-decoding results in our simulated data, demonstrating that it is highly unlikely that our pattern of cross-decoding results can be accounted for by differences in signal-to-noise ratio. Our simulation analysis thus converged on the interpretation that preparatory overt attention recruits an additional spatially selective process that is absent during covert shifts of attention.

Together, our analyses demonstrate preparatory overt attention to recruit (at least) one additional spatially selective neural process that was not present when covert attention was shifted. This additional spatial selectivity was most prominent in frontal channels, and explains the sharper spatial selectivity in preparatory overt than in covert attention. We speculate that motor preparation and/or predictive remapping underlie this additional spatially selective process (see Discussion).

## Discussion

We here demonstrated that population-level activity as reflected in EEG (Lopes da Silva, 2013; Nunez, 2006) dissociates preparatory overt from covert attention. This dissociation emerged rapidly after cue onset, and was in part driven by the recruitment of (at least) one additional spatially selective process during preparatory overt attention. Our results dissociate preparatory overt from covert attention at a much broader scale than previously proposed.

By definition the neural signatures of preparatory overt and covert attention must diverge as a saccade is initiated in the former but not the latter case (Awh et al., 2006). But when do these neural signatures diverge: Does this already occur prior to saccade initiation? Complementing evidence from single-cell recordings (Messinger et al., 2021), our decoding results demonstrate that population-level neural activity dissociated rapidly after cue onset. Moreover, we found that spatially selective neural activity emerged earlier during preparatory overt compared with covert attention. These data resonate with psychophysical data demonstrating differential effects of presaccadic and (voluntary) covert attention on perception rapidly after cue onset (Li, Hanning, & Carrasco, 2021; Li, Pan, & Carrasco, 2021; Rolfs & Carrasco, 2012). Our results reveal a rapid neural dissociation between preparatory overt and covert attention well before saccade onset.

We previously found a larger pupil size during preparatory overt attention compared with covert attentional shifts (Koevoet et al., 2023) - which we replicated here - and hypothesized that this larger pupil size could be attributed to the recruitment of additional neural processes in preparatory overt attention. We here found direct neural evidence for this hypothesis, as cross-decoding revealed an additional spatially selective process in frontal cortex in preparatory overt attention (Figure 5; Figure 6). What could this additional spatially selective process reflect?

Although our data cannot cannot answer this question with certitude, we here discuss two possible candidates that are inherently spatial and recruit FEF: Motor preparation and predictive remapping (Awh et al., 2006; Bruce et al., 1985; Corbetta, 1998; Deubel & Schneider, 1996; Duhamel et al., 1992; Jonikaitis & Moore, 2019; Kowler et al., 1995; Li, Hanning, & Carrasco, 2021; Mayo et al., 2015; Messinger et al., 2021; Moore & Fallah, 2001; Moore et al., 2003; Perry & Zeki, 2000; Rao et al., 2016; Rizzolatti et al., 1987, 1994; Schall, 2015; Sparks, 2002). Jonikaitis et al. (2023) demonstrated that pharmacological inactivation (using muscinol) of FEF impaired motor preparation but not spatial memory of the target location (or covert spatial attention, see Awh & Jonides, 2001; Awh et al., 1998; Foster, Bsales, et al., 2017; Foster et al., 2016; Williams et al., 2013). In line with this, single-unit recordings in FEF revealed enhanced spatial selectivity when a saccade can be prepared to the target compared with solely attending/memorizing the target location (Park et al., 2025). As for the timecourse, the independent contribution of motor preparation to spatial selectivity emerges well before saccadic onset, which closely fits with our population-level results (Messinger et al., 2021; Park et al., 2025) (also see Gutteling et al., 2010). The above puts forth motor preparation as a strong candidate to underlie the additional spatially selective process we observed in frontal cortex during preparatory overt but not covert attention.

Another candidate that could underlie the additional spatially selectivity is predictive remapping. Predictive remapping entails the remapping of receptive fields from the presaccadic stimulus location to the post-saccadic stimulus location. Such remapping accounts for displacements of the visual world on the retina due to saccades, which facilitates visual continuity (Cavanagh et al., 2010; Currie et al., 2000; Deubel & Schneider, 1994; Deubel et al., 1998; Fabius et al., 2019; Koevoet, Naber, et al., 2025; Mathôt & Theeuwes, 2011; McConkie & Currie, 1996; Rolfs, 2015; Szinte et al., 2018; Van der Stigchel & Hollingworth, 2018). Importantly, predictive remapping should therefore only occur in preparatory overt but not covert attention (Duhamel et al., 1992; Smith & Schenk, 2012; Van der Stigchel & Hollingworth, 2018). Although the above argues for a potential role of predictive remapping in explaining our results, the timecourse of predictive remapping may not fit our findings. For example, recent M/EEG decoding studies demonstrated that the post-saccadic stimulus could be decoded only briefly (∼50-100 ms) preceding saccade onset (Fabius et al., 2020; Moran et al., 2024), and similar timecourses are observed using single-unit FEF recordings (e.g. Wang et al., 2024). By contrast, in the current data preparatory overt attention already showed sharper spatial selectivity than covert attention well before saccade initiation (>550 ms). One discrepancy is that participants initiated saccades as fast as possible upon cue onset in previous work (Fabius et al., 2020; Moran et al., 2024) but delayed their saccade in our task, making it difficult to compare timecourses between them. Future work should address whether preparatory overt also shows sharper spatial selectivity than covert attention when there is no delay (i.e. saccades are initiated as fast as possible upon cue onset). Note that we are unsure whether participants even *could* predictively remap array stimuli since they were not presented prior to saccade onset. Nevertheless, we cannot rule out that the fixation point was remapped predictively. Thus, predictive remapping may account for the sharper spatial tuning involved in preparatory overt than covert attention but its timecourse may not be compatible with the current results. We speculate that the additional frontal spatially selective process likely reflects motor preparation as previous work provides more traction for this than predictive remapping but more work is necessary to definitively arbitrate between these candidates.

Where do our results leave motor-based theories of visual attention? For example, the popular premotor theory of attention (Rizzolatti et al., 1987, 1994) equates covert attention to preparing but not executing a saccade. Our results oppose a ‘hard’ interpretation of the premotor theory since we demonstrate dissociable neural activity between preparatory overt and covert attention. Nevertheless, we did observe cross-decodability of cued locations across conditions, demonstrating overlap in attentional mechanisms. One way that a weaker interpretation of the premotor theory may still hold, is if the additional spatially selective process observed here reflects predictive remapping (Koevoet et al., 2023; Smith & Schenk, 2012). In such a view, preparatory overt attention is tantamount to covert attention but with an added spatially selective non-motor process. However, if the observed additional spatially selective process reflects motor preparation, our data would show that this weaker version of the premotor theory of attention breaks down even at the population level.

Throughout the manuscript we created topographies of our decoding analyses using a searchlight approach (Kriegeskorte et al., 2006). While multivariate classification revealed that preparatory overt and covert attention could be discerned across the scalp (Figure 3), IEM analyses revealed that particularly frontal channels contributed to differences in decoding (Figure 6). Using the IEMs we were able to detail differences in spatial selectivity between preparatory overt and covert attention. However, the whole-scalp contribution to multivariate classification suggests that other, space-agnostic processes may also differ between preparatory overt and covert attention. Among other options, it is possible that non-spatial action-readiness signals or underlying perceptual computations of preparatory overt and covert attention (Li, Pan, & Carrasco, 2021) are neurally represented in a non-spatial format. Such non-spatial signals account for the different topographies obtained with the multivariate classification and IEM analyses. Thus, the searchlight topographies indicate that preparatory overt and covert attention may differ beyond spatial signals, yet the potential functions of such non-spatial activity remain open.

In contrast to past work using similar tasks that found no differences (e.g. Gutteling et al., 2010; Koevoet et al., 2023), we found higher accuracy in preparatory overt compared with covert trials. We here speculate what may account for this accuracy difference. We found that the accuracy difference could not be predicted by pupil size, nor by differences in CTF slopes across participants. However, due to overall relatively high accuracy on the task, we did not have access to many incorrect trials, and were unable to directly compare correct with incorrect trials within participants. It is therefore possible that with more trials or when using more difficult tasks with overall lower accuracies that CTF slopes may predict task accuracy. Another possibility is that the additional frontal spatially selective process in preparatory overt attention does not contribute to perception, but instead reflects other processes (such as motor preparation). In that case, differences in accuracy may be accounted for by differential underlying computations of presaccadic and voluntary covert attention (Li, Pan, & Carrasco, 2021). Nevertheless, more work is necessary to examine what may drive accuracy differences between preparatory overt and covert attention, as well as its possible neural correlates.

Together, we extend previous dissociations between preparatory overt and covert attention from single cells to the population level. We found that preparatory overt and covert attention dissociated rapidly. Specifically, an additional spatially selective process in frontal cortex was recruited in preparatory overt but not in covert attention. We conclude that population-level activity dissociates preparatory overt from covert attention at a much larger scale than previously thought.

## Acknowledgments

This project has received funding from the European Research Council (ERC) under the European Union’s Horizon 2020 research and innovation programme (grant agreement n° 863732). A part of this work was presented during the Vision Sciences Society 2025 meeting (Koevoet, Voet, et al., 2025).

## Footnotes

1 We use the label ‘preparatory overt attention’ to stress the fact that no saccade has yet been executed. Note that we do not use ‘saccade preparation’, as premotor theories of attention posit that covert spatial attention also involves saccade preparation. We therefore chose ‘preparatory overt’ as a theoretically neutral label.

2 We also examined whether the pupil effect between conditions (*β*) predicted the accuracy difference between the preparatory overt and covert conditions across participants. We observed no significant correlation (*r*(21) = -.013, *p* = .95).

