## Supplementary Information for "Population-Level Activity Dissociates Preparatory Overt from Covert Attention"

### Supplementary Text 1: Alpha Oscillations Track Spatial Attention

Although we focused on analyzing voltage data in the main paper, inverted encoding modeling has popularly been applied to alpha oscillations (Foster et al., 2016, 2017, 2020; Samaha et al., 2016; van Moorselaar et al., 2018). For completeness, we here also present results when using inverted encoding modeling using alpha power. We computed alpha power using the filter-Hilbert transform method (Cohen, 2014). Specifically, we first bandpass filtered the data to isolate activity from the alpha-band (8-12 Hz). We then applied a Hilbert transform to obtain the complex analytical signal which we subsequently squared to compute alpha power at each timepoint. This array then served as input to the inverted encoding models.

In line with previous work, we observed spatially selective alpha power when participants shifted covert attention (156-700 ms, $p < .0001$). Moreover, we observed a similar pattern in preparatory overt attention was shifted (227-700 ms, $p < .0001$). This shows that alpha power reliably tracked the attended location. Alpha power also showed a slightly sharper spatially-tuned activity to the attended location in preparatory overt compared with covert attention (320-469 ms, $p = .039$). This cluster was centered around 400 ms, and seemed to differ from the voltage analysis in terms of timing. In the voltage analysis, the difference between conditions was strongest within earlier and later windows.

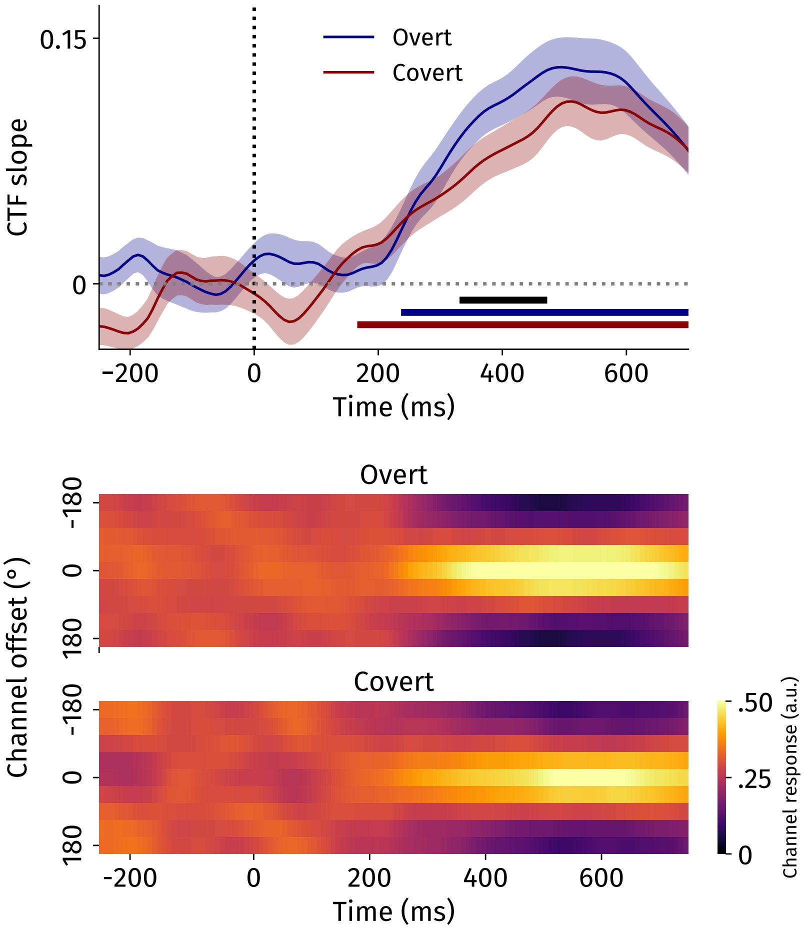


Supplementary Figure 1: Alpha power-based inverted-encoding modeling results. Horizontal blue and red lines represent clusters where CTF slopes are significantly above 0 ($p < .0001$) for the overt and covert data, respectively. Horizontal black line indicates a significant cluster between conditions ($p < .05$). Error bands reflect standard errors of the mean. The lower plot shows heatmaps separately per shift type to visualize channel responses from the reconstructed CTFs across time.

### Supplementary Text 2: Eye-movement Control Analyses

#### Multivariate Classification Control Analysis

In the main paper, we could reliably decode whether participants shifted preparatory overt or covert attention rapidly after cue onset. We here wanted to rule out that this effect was driven by eye movements. To this end, we first reran the classifier using only drift-corrected (-250 until 0 around cue onset) gaze data (Supplementary Figure 2A,B). We could reliably classify trials based on the gaze data (even slightly before cue onset).

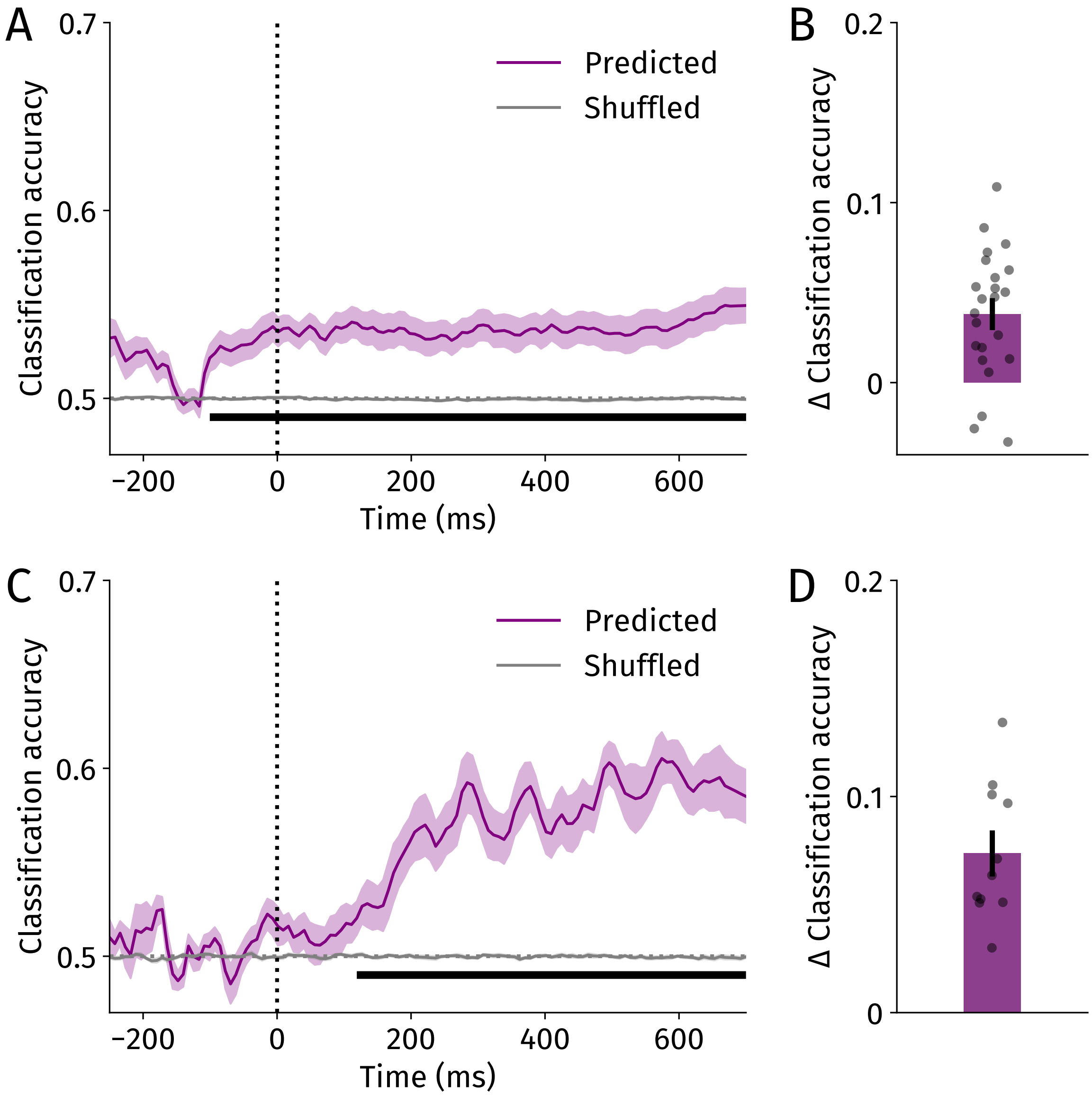


Supplementary Figure 2: Multivariate classification eye movement control analyses. **A** Time-resolved classification of shift type (overt or covert) using only gaze data. **B** Average (100-700 ms) difference in classification accuracy difference between the actual and empirical chance based on gaze data. **C** Time-resolved classification of shift type (overt or covert) using neural data of a subset of ten participants without informative gaze. **D** Average (100-700 ms) difference in classification accuracy difference between the actual and empirical chance based on EEG data for the subset of participants without informative gaze. In **A, C**, horizontal black lines indicate significant clusters ($p < .05$). In **B, D**, black dots reflect individual participants. Error bands and bars reflect standard errors of the mean.

To rule out that our neural classification was caused by gaze patterns, we selected a subset of participants that had a relatively low classification accuracy based on gaze (< 0.04 ΔClassification accuracy averaged 100-700 ms post-cue) (Supplementary Figure 2C,D). In this subset of eleven participants, trials could no longer be classified reliably using gaze patterns (all cluster *p*s > .10). In this subset of ten participants without informative gaze, we could still reliably decode whether they shifted preparatory overt or covert attention based on their EEG data (109-700 ms, $p < .001$). This rules out eye movements as an important confound to our classification results.

### Inverted Encoding Modeling Control Analysis

To ascertain whether gaze contributed to our inverted-encoding modeling (IEM) results, we first tested whether miniature gaze patterns were spatially selective to the cued location at all. We calculated the distance of gaze from the cued location separately for the overt and covert conditions (Supplementary Figure 3). To account for drift, we baseline-corrected the gaze data -250 - 0 around cue onset. We observed very small gaze patterns (<0.1°) toward the cued location in both conditions (overt: 219-700 ms, $p < .0001$; covert: 215-700 ms, $p < .0001$). Note that the onset of the gaze bias was around 215 ms, therefore these pattern cannot account for our earliest cluster (starting 144 ms after cue) found in the main IEM analysis. As for the later window, we found that gaze was significantly more spatially selective in the overt compared with the covert condition (382-700 ms, $p = .015$).

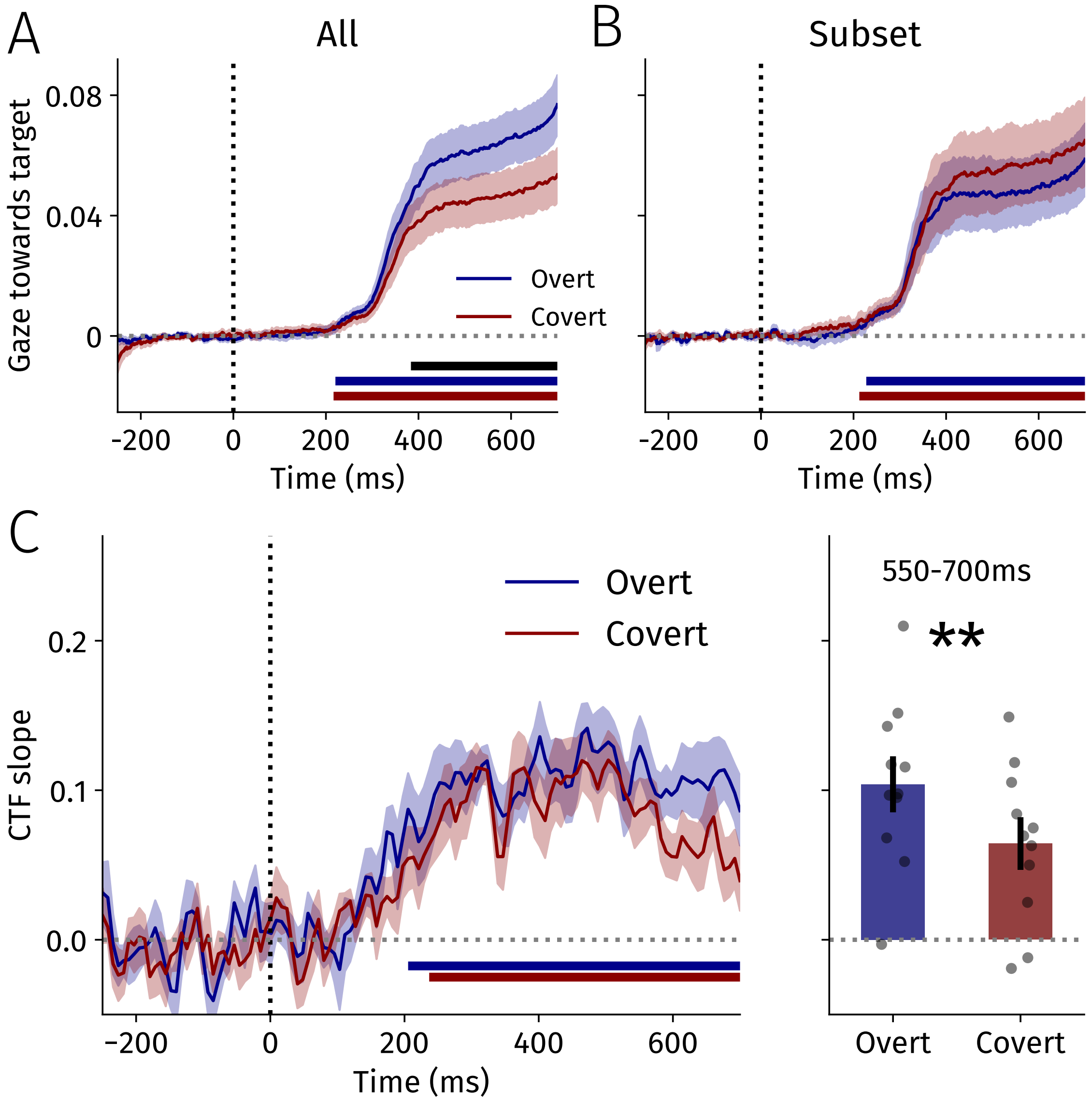


Supplementary Figure 3: **A** Spatial selectivity of gaze for all participants. **B** Spatial selectivity of gaze for the subset of participants. **C** Inverted encoding modeling results for the gaze-selected subset. We averaged CTF slopes across the late window and compared them across conditions. Horizontal blue and red lines represent clusters of significant spatial selectivity of gaze (**A**, **B**) and CTF slopes (**C**) ($p < .0001$) for the overt and covert data, respectively. Horizontal black bar indicates significant difference between conditions ($p < .05$). Error bands reflect standard errors of the mean.

We wanted to test whether higher neural spatial selectivity in the overt compared with covert condition in a later time window (550-700 ms after cue onset) could be ascribed to eye movements. As a crucial test to see whether sharper tuning in the overt condition was caused by more spatially selective gaze, we used a median-split to select participants that contributed the least to the difference between overt and covert in spatial selectivity of gaze (Supplementary Figure 3B). We then reconducted the IEM analysis on these eleven participants. We reasoned that if in these participants we would still find the difference in the expected direction, differences in tuning in the later time window could not be attributed to eye movements.

In this subset of eleven participants (Supplementary Figure 3C), we found spatially-tuned neural responses to the cued location in both the overt and covert conditions (overt: 203-700 ms, $p < .001$, covert: 234-700 ms, $p = .003$). We averaged CTF slopes across the later timewindow (550-700 ms) and compared conditions using a two-tailed paired-samples *t*-test. Consistent with the analysis reported in the main paper, we found sharper spatially-tuned neural responses in the overt compared with the covert condition ($t(10) = 3.46$, $p = .006$, $d = 1.04$). Thus, the observed sharper spatially-tuned neural responses in preparatory overt compared with covert attention is driven by neural activity and not by gaze patterns.

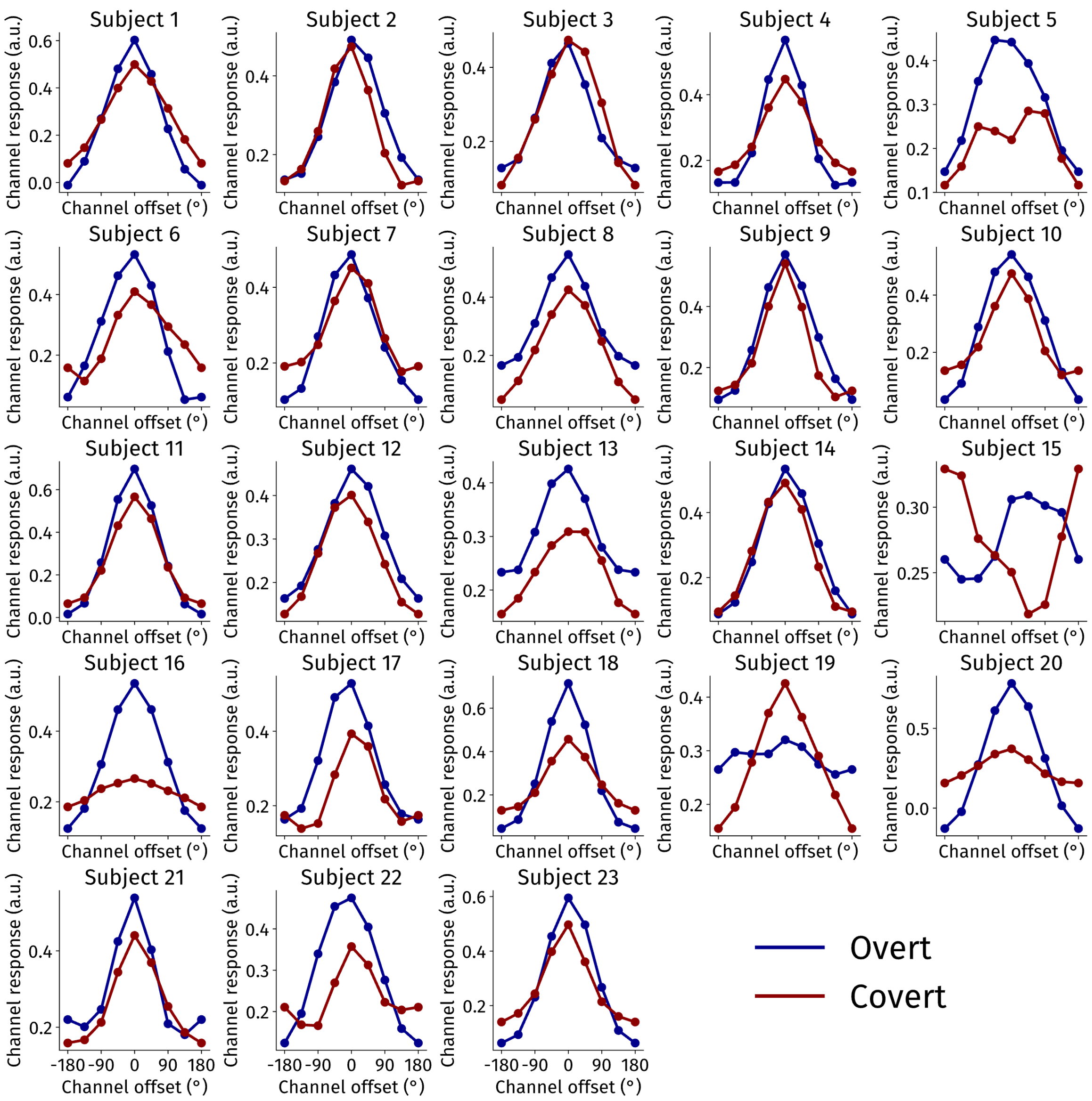


Supplementary Figure 4: Reconstructed CTFs for each participant separately. Specifically, we averaged across the delay period (100-700 ms) for each participant and plot reconstructed CTFs (recentered and meaned across locations).

### Supplementary Text 3: Spatial Selectivity Weakly Modulates Response Times

To test whether the spatial selectivity of neural activity was predictive of response times, we conducted an exploratory analysis. Specifically, we used median splits separately per shift type condition. By splitting separately per condition, potential differences in spatial selectivity in fast or slow response time trials could not be attributed to shift type differences. We then conducted our IEM analysis as in the main paper.

For both fast and slow response time trials, we found substantial spatially selective neural activity to the target location ($ps < .0001$). For a brief moment, we observed more spatially selective neural activity in fast compared with slow response time trials (273-315 ms; $ps = .007$). This effect was not significant when averaging across the delay period ($t(22) = 2.06$, $p = .052$, $d = 0.43$). We therefore conclude that spatial selectivity only weakly modulates the speed of the ultimate response at the end of the trial. The effect in the current data is relatively weak compared with previous work by Foster et al. (2017). We speculate that this is likely because we here employed a cued selection task, while Foster et al. (2017) used visual search task without any cues. In that case, spatially selective neural activity should more closely be linked to response times, as in that task participants respond as soon as the target is identified. Moreover, the latency differences that were tracked in the Foster et al. (2017) study were much larger than the cueing effects in the present dataset.

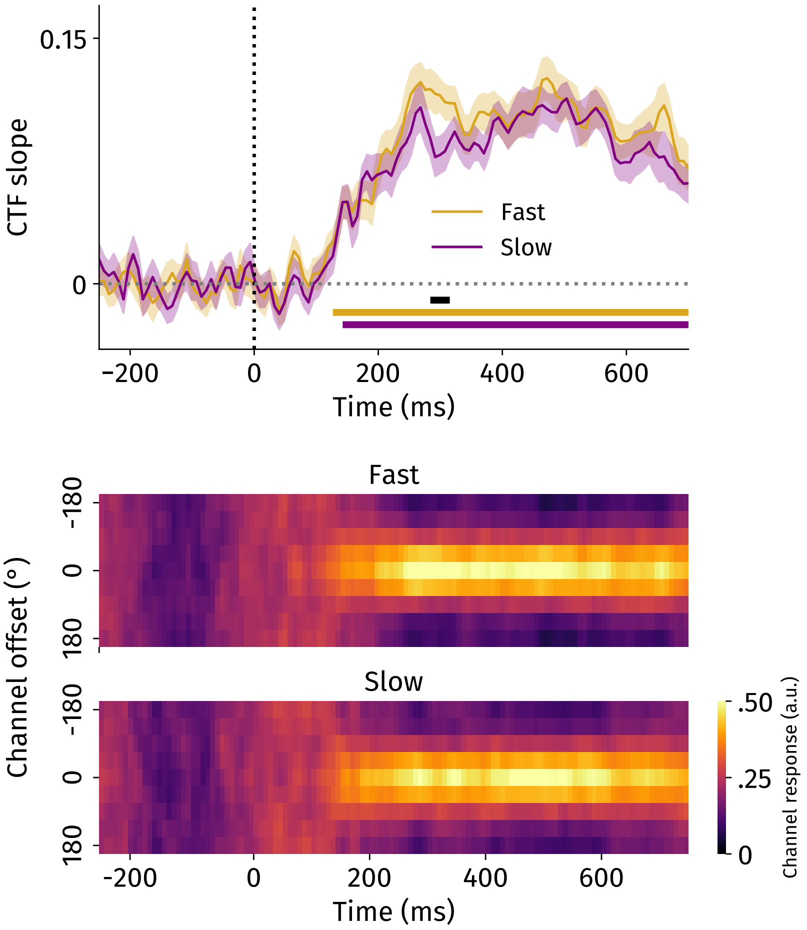


Supplementary Figure 5: Spatial selectivity of as a function of response speed. Horizontal orange and purple lines represent clusters of spatial selectivity ($p < .0001$) for fast and slow response data, respectively. Horizontal black bar indicates significant difference between condition ($p < .05$). Error bands reflect standard errors of the mean.

### Supplementary Text 4: Ruling Out Differences in Signal-To-Noise Ratio as an Alternative Explanation

Past work suggests that, in some cases, cross-decoding asymmetries may driven by signal-to-noise ratio (SNR) instead of neurocognitive effects. As one of our main findings is most strongly supported by cross-decoding results, we wanted to examine whether SNR differences could reproduce our observed effects. Here, we first describe the results from simulation work by van den Hurk and Op de Beeck (2019). We then provide arguments why it is unlikely that our results were driven by differences in SNR between conditions. Lastly, we created our own simulations based on properties of our observed EEG data to investigate whether our findings were driven by SNR.

van den Hurk and Op de Beeck (2019) investigated whether asymmetries in cross-decoding could be driven by differences in SNR between conditions. Cross-decoding asymmetries refer to the phenomenon that when training on condition A, decoding accuracy is higher than when training solely on condition B. This is akin to our cross-decoding IEM results. Using simulations, the authors demonstrated that when there are substantial differences in SNR between conditions, cross-decoding asymmetries can occur. That is, asymmetries occur because of SNR differences and not because of actual differences between conditions.

We deem it unlikely that our results were driven by differences in SNR for three reasons. First, SNR-driven decoding asymmetries occur when training on the 'noisy' condition and then tested on the higher SNR condition. However, as CTF slopes are overall steeper in the preparatory overt than in the covert attention condition, if SNR drove our effects, it should be higher in preparatory overt than in covert attention trials. Following this, our cross-decoding results actually oppose simulated SNR-driven decoding asymmetries, strongly arguing for a true cross-decoding asymmetry caused by actual neurocognitive effects. Second, SNR only drives cross-decoding asymmetries when there are substantial differences in within-condition decoding accuracy between the conditions. In their example, this occurs when binary decoding accuracy differed by $\sim$10% between conditions. We did find differences in within-condition decoding CTF slopes, but these did not seem as substantial as in the example – although it is difficult to compare classification accuracies and CTF slopes directly. Third, searchlight analyses demonstrated that preparatory overt attention recruits an additional spatially selective signal in frontal regions (Figure 6). An SNR account would not predict such distinct topographies.

To complement these arguments, we created simulations to assess this quantitatively. Simulations were performed in the following steps. First, we fit the IEM across both shift type blocks on the trial level across 10 iterations per participant. Note that for each iteration, we ensured an equal number of trials per unique conditions (2 shift types x 8 locations). This allowed us to obtain an average weights matrix per participant. Moreover, during each iteration, we also calculated the covariance matrix (using the Ledoit-Wolff procedure). Second, in order to simulate data, for each participant we used the averaged weights matrix across iterations and the basis channel responses to simulate spatially selective data. Third, we applied scalars to these data separately per shift type block to scale their spatially selective signals. In order to recreate data that resembled our original results (from Figure 4A), we chose scalars 1.5 and 0.75 for the preparatory overt and covert conditions, respectively. Fourth, after scaling the data, we added noise from a multivariate normal distribution with a mean vector of 0 at each electrode, and a covariance structure equal to the average covariance matrix computed across the 10 iterations above. As we used these covariance matrices, the noise in the data resembled the original EEG data, making our simulations more comparable to our original analyses. Fifth, using these data, we conducted the same IEM analysis as in the main paper, as well as the cross-decoding results. Due to overfitting (see below), we observed significant spatial selectivity in the baseline period, and we therefore baseline-corrected the obtained CTF slopes.

We were able to recreate the results from Figure 4A in the main paper using our simulated data (Supplementary Figure 6A). As intended, we found sharper spatial selectivity in the simulated preparatory overt compared with the simulated covert condition (averaged across delay: $t(22) = 5.83$, $p < .001$, $d = 1.24$). Crucially, in our simulated data, all differences between conditions were purely driven by differences in SNR. This allowed us to examine whether our reported cross-decoding results were driven by SNR.

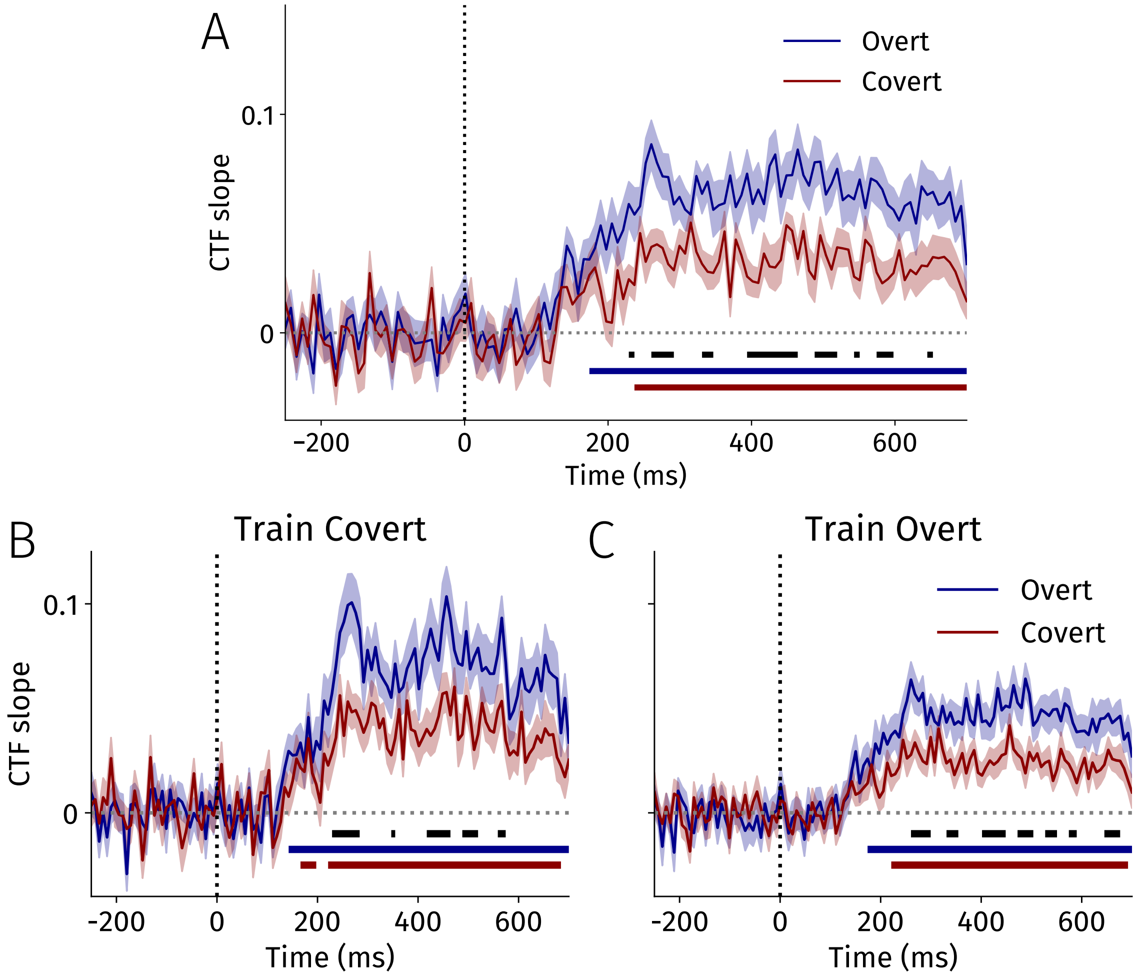


Supplementary Figure 6: **A** Simulation-based inverted-encoding modeling results when training on both conditions. **B, C** Simulation-based cross-decoding results. Horizontal blue and red lines represent clusters where CTF slopes are significantly above 0 ($p$ < .05) for the overt and covert data, respectively. Horizontal black line indicates a significant cluster between conditions ($p$ < .05). Error bands reflect standard errors of the mean.

The cross-decoding results revealed a different pattern than that reported in our main paper (Supplementary Figure 6B,C). Specifically, regardless of the training condition, we observed sharper spatial selectivity in the simulated preparatory overt compared with the simulated covert condition (averaged across delay, train covert: $t(22) = 6.18$, $p < .001$, $d = 1.32$; train overt: $t(22) = 6.09$, $p < .001$, $d = 1.30$). Moreover, averaging across the delay period (100-700ms postcue), we found evidence for a modest interaction effect ($t(22) = 2.08$, $p = .049$, $d = .44$). That is, when training on the covert data, differences in spatial selectivity between simulated preparatory overt and simulated covert data were larger when compared to training exclusively on the preparatory overt data. This simulation demonstrates that, if effects are purely driven by SNR, cross-decoding results would strongly oppose the additional source account. Therefore, these simulation results strongly suggest that our main cross-decoding results do not reflect differences in SNR but instead reveal a neurocognitive effect.

Our simulation analyses had some limitations. First, some aspects of our simulation results differ slightly when adjusting the scalar values. For example, in some cases we did not observe an interaction effect, but only observed preparatory overt to show sharper spatial selectivity regardless of training condition (i.e. a main effect). Our conclusions do not hinge on this interaction effect, however, as even in that case we could not recreate the cross-decoding results reported in the main paper by solely manipulating SNR. Second, because of noise in the baseline period, the IEM weights computed during the baseline were always non-zero, i.e. overfit. As a result, we believe our simulation procedure generated reliable patterns of activity even during the baseline, leading to significant spatial selectivity prior to the cue onset. In support of this overfitting explanation, we found that the weights during the baseline were highly unstable and variable across adjacent time points, while the weights obtained during post-cue timepoints were highly reliable across time. To correct for this overfitting, we baseline-corrected the obtained CTF slopes in all of the analyses. Note that this interacted with the scalars that we chose in order to reproduce Figure 4A, as changing the scale also changes the selectivity in the baseline period. As noted above, while different scalar choices impact the strength of the interaction, we could never qualitatively reproduce the cross-decoding results we see in our real data via manipulations of SNR. While out of scope for the current purpose, future work could address the above limitations by using cross-validation throughout the entire simulation process, and by creating simulations across many different combinations of scalar values.

Despite these limitations, however, our simulation was able to recover one core finding from van den Hurk and Op de Beeck (2019) as training exclusively on the low SNR condition (covert) yielded overall sharper spatial selectivity compared with training exclusively on the high SNR condition (overt). Most importantly, we here report an example where simulated data (based on properties of the observed EEG data) that only differed in SNR yielded very different results when cross-decoding. Together, based on prior simulation work, the differences in topographic maps based on training and testing conditions (Figure 6), as well as the current simulations incorporating properties of the observed EEG data, we deem it highly unlikely that SNR can account for our reported cross-decoding asymmetry.
